# Uncovering the molecular interactions underlying MBD2 and MBD3 phase separation

**DOI:** 10.1101/2024.04.29.591564

**Authors:** Nicole Maurici, Tien M. Phan, Jessica L. Henty-Ridilla, Young C. Kim, Jeetain Mittal, Alaji Bah

**Affiliations:** Department of Biochemistry and Molecular Biology, SUNY Upstate Medical University, Syracuse, NY, 13210; Artie McFerrin Department of Chemical Engineering, Texas A&M University, College Station, TX, USA; Department of Neuroscience and Physiology, SUNY Upstate Medical University, Syracuse, NY, 13210; Center for Materials Physics and Technology, Naval Research Laboratory, Washington, DC, USA; Department of Chemistry, Texas A&M University, College Station, TX, USA; Interdisciplinary Graduate Program in Genetics and Genomics, Texas A&M University, College Station, TX, USA

## Abstract

Chromatin organization controls DNA’s accessibility to regulatory factors to influence gene expression. Heterochromatin, or transcriptionally silent chromatin enriched in methylated DNA and methylated histone tails, self-assembles through multivalent interactions with its associated proteins into a condensed, but dynamic state. Liquid-liquid phase separation (LLPS) of key heterochromatin regulators, such as heterochromatin protein 1 (HP1), plays an essential role in heterochromatin assembly and function. Methyl-CpG-binding protein 2 (MeCP2), the most studied member of the methyl-CpG-binding domain (MBD) family of proteins, has been recently shown to undergo LLPS in the absence and presence of methylated DNA. These studies provide a new mechanistic framework for understanding the role of methylated DNA and its readers in heterochromatin formation. However, the details of the molecular interactions by which other MBD family members undergo LLPS to mediate genome organization and transcriptional regulation are not fully understood. Here, we focus on two MBD proteins, MBD2 and MBD3, that have distinct but interdependent roles in gene regulation. Using an integrated computational and experimental approach, we uncover the homotypic and heterotypic interactions governing MBD2 and MBD3 phase separation and DNA’s influence on this process. We show that despite sharing the highest sequence identity and structural homology among all the MBD protein family members, MBD2 and MBD3 exhibit differing residue patterns resulting in distinct phase separation mechanisms. Understanding the molecular underpinnings of MBD protein condensation offers insights into the higher-order, LLPS-mediated organization of heterochromatin.

## INTRODUCTION

The nucleus, being densely packed, is an ideal environment for self-organizing membraneless bodies and contains liquid-liquid phase separation (LLPS)-regulated membraneless organelles (MLOs) that include the nucleolus, Cajal bodies, paraspeckles, and transcription factories (1, 2). Recent studies suggest LLPS plays a part in heterochromatin formation and regulation, resulting in the formation of spatiotemporally regulated MLOs composed of genomic DNA and its associated proteins (3–6). The assembly and dissolution of these heterochromatin condensates contribute to genome organization and regulation of gene transcription (3, 7–9). Specifically, proteins that bind to modified histones and methylated DNA communicate between DNA methylation- and histone modification-associated complexes to carry out heterochromatin condensation through LLPS (5, 9, 10). While phase separation of histone-associated proteins and their role in heterochromatin formation and function have been investigated in molecular detail, phase separation of methyl-binding proteins and their influence on heterochromatin are just beginning to be studied (11, 12).

Methyl-binding proteins include the methyl-CpG-binding domain (MBD) family of proteins that are localized at methylated CpG islands near gene promoter regions and pericentric heterochromatin (chromocenters) where they bind to methylated CpG dinucleotides (13–16). As transcriptional repressors, they are responsible for regulating the transcriptional state of the genome by recruiting and coordinating the interactions, or crosstalk, of macromolecular systems responsible for DNA methylation, histone modification, and chromatin organization. Previous studies indicate that LLPS of heterochromatin protein 1α (HP1α) that binds to methylated histone tails plays an essential role in heterochromatin assembly (3, 6, 8, 17). HP1α is also known to interact with several MBD proteins and co-phase separates with the well-characterized MBD protein MeCP2 (5, 10, 18). MeCP2 has been recently shown to undergo LLPS with and without methylated DNA, and, crucially, mutations within MeCP2 known to cause Rett syndrome result in aberrant condensate formation, highlighting the importance of LLPS in disease pathology (5, 9, 19). While these studies provided an initial foundational understanding, the molecular mechanisms by which MeCP2 undergoes LLPS remain to be fully elucidated. Additionally, it is not clear if other members of the MBD family share this ability to undergo LLPS to mediate their functions in influencing genome organization and transcriptional regulation.

To further our understanding of the role of methyl-binding protein mediated-LLPS in heterochromatin assembly, we have chosen to focus on two MBD proteins, MBD2 and MBD3. Like MeCP2, and similar to HP1α, MBD2 and MBD3 are modular in their domain architecture containing both folded domains and large, compositionally biased intrinsically disordered regions (IDRs) important in generating transient, multivalent interactions that promote LLPS (7–9, 15, 20) (Figure 1A and B). Interestingly, MBD2 and MBD3 share the highest sequence similarity (71.1%) among all members of the MBD family. They both possess an MBD and coiled-coil (CC) domain flanked by IDRs: however, their compositions vary in terms of their IDRs and the presence or absence of specific domains. Most notably, MBD2 contains a transcriptional repression domain (TRD), and its N-terminus is enriched in arginine and glycine residues while MBD3’s C-terminus contains an aspartic/glutamic-rich region (Figures 1A and B). Of importance to note, MBD3, although it contains an MBD, does not bind to methylated DNA with high affinity due to two amino acid substitutions (15, 21–24). MBD2 and MBD3 have roles in heterochromatin formation and transcriptional repression, like MeCP2. They are both an integral part of the Nucleosome Remodeling and Deacetylase (NuRD) complex but differ in their spatiotemporal expression patterns and where they bind on the genome as demonstrated by genetic knockout and knockin experiments in mice (25). Because of these differential genomic binding profiles and expression patterns, they also have their own respective roles. Despite this, their activities seem to be regulated by each other (23, 25–29). Therefore, we wanted to explore the mechanisms underlying MBD2 and MBD3 LLPS as the functional roles of their IDRs and their ability to undergo LLPS have yet to be explored (15, 30, 31). Their intrinsic disorder has posed challenges for biophysical studies thus far, primarily due to poor protein yields. However, we have optimized expression and purification methods tailored specifically for intrinsically disordered proteins (IDPs), enabling us to produce sufficient quantities of full-length (FL) protein for *in vitro* characterization (32). Determining how MBD2 and MBD3 are involved in condensate formation will be important for understanding how aberrations in their ability to interact with themselves and other binding partners lead to dysregulation in condensate dynamics and, consequently, their roles in heterochromatin formation and transcriptional repression.

**Figure 1.**
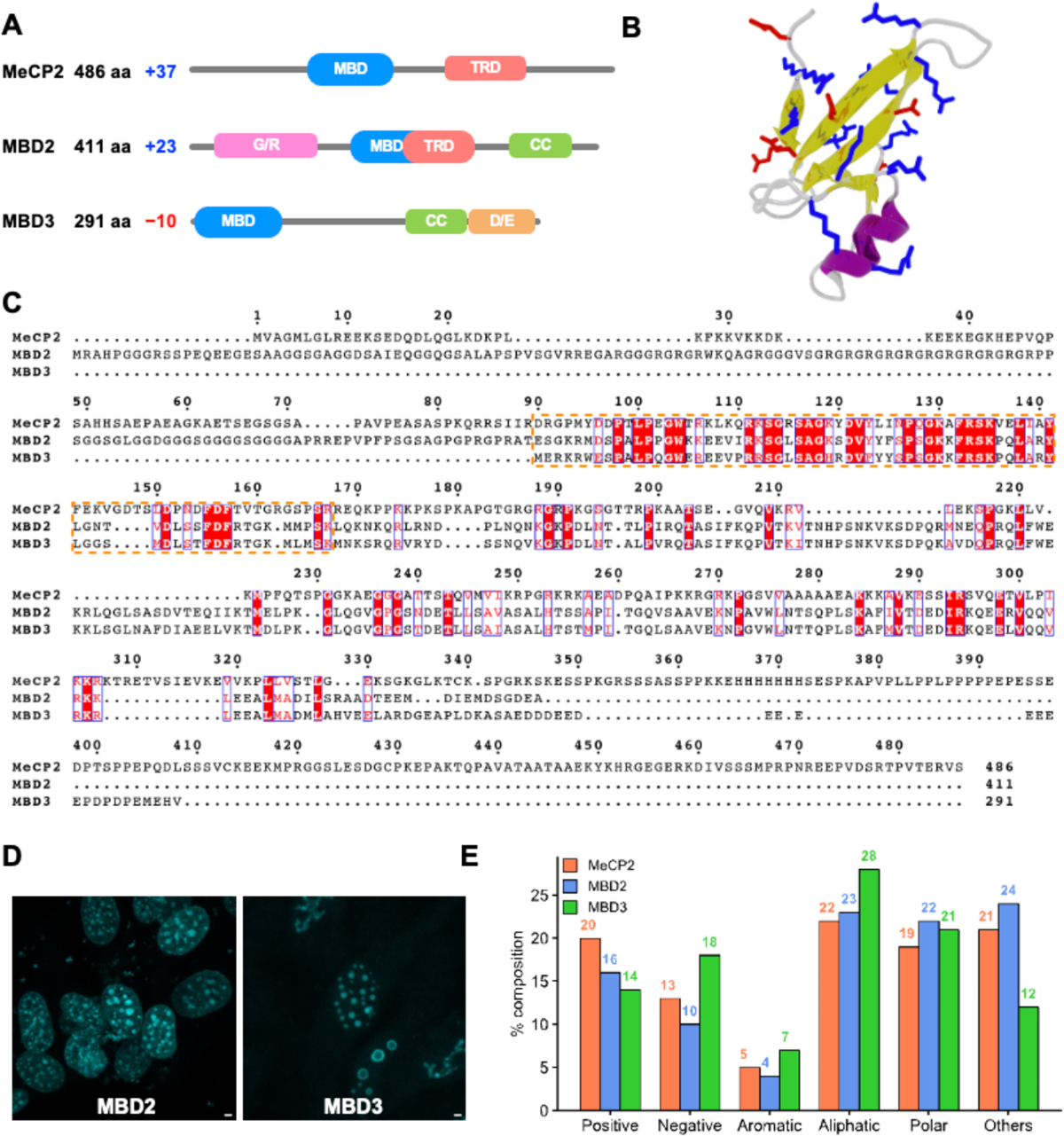
Properties of MBD proteins. **A.** Protein domain architecture maps of MeCP2, MBD2, and MBD3. The domains represented are MBD: methyl binding domain, TRD: transcription repression domain, G/R: glycine/arginine-rich domain, CC: coiled-coil domain, and D/E: aspartic/glutamic rich domain. **B.** Structure of the methyl-CpG-binding domain (PDB ID: 2mb7). Positively and negatively charged residues are shown in blue and red licorice representation, respectively. **C.** Multiple sequence alignment of MeCP2, MBD2, and MBD3 using Clustal Omega and ESPript web servers (54, 55). Red boxes with white letters show identical amino acids, and white boxes with red letters indicate amino acids with similar properties. The orange box highlights the conserved methyl-CpG-binding domain. **D.** MBD2 and MBD3 form condensates *in cellulo*. Fixed 3T3 fibroblast cells imaged by laser scanning confocal microscopy illuminating nuclear condensates of eGFP-tagged, FL MBD2 (left) or FL MBD3 (right). Scale bar = 2 µm. **E.** Amino acid composition of MeCP2, MBD2, and MBD3 are organized into positive (Arg, Lys), negative (Asp, Glu), aromatic (His, Phe, Tyr, Trp), aliphatic (Ala, Ile, Leu, Met, Val), polar (Asn, Gln, Ser, Thr), and others (Cys, Gly, Pro).

Herein, we employ a variety of computational and experimental techniques to assess MBD2 and MBD3’s ability to phase separate and the driving forces that allow them to do so. We have identified regions and sequence features within MBD2 and MBD3 responsible for driving their LLPS and have demonstrated their propensity to phase separate individually, together, and with and without methylated DNA *in vitro* and *in cellulo*. We have determined the conditions that induce their characteristic LLPS behavior and show how interactions with each other and methylated DNA influence their LLPS propensities. These studies provide a framework for understanding how the interactions within MBD-containing nucleoprotein complexes mediate LLPS to regulate chromatin condensation and transcription.

## MATERIALS AND METHODS

### Expression and purification of MBD2 and MBD3 and their respective truncations

Codon-optimized human cDNA of FL and truncated MBD2 and MBD3 were cloned into a pET28 vector containing an N-terminal His_6_-SUMO tag, and kanamycin resistance (33). The cysteines within the proteins were mutated to serines by site-directed mutagenesis to prevent non-native disulfide bond formation and reduce aggregation. The plasmids encoding the proteins were transformed into chemically competent BL21-CodonPlus (DE3) RIPL *E. coli* cells and grown in LB media containing 50 μg mL^-1^ kanamycin and 34 μg mL^-1^ chloramphenicol to an optical density (OD_600_) of ∼0.6-1.0 at 37°C. The cultures were induced with 1 mM IPTG at 16°C and harvested after 12-18 hours by centrifugation at 4, 000 RPM at 4°C for 30 minutes. Finally, the cells were resuspended with lysis buffer (300 mM NaCl, 50 mM Na_2_PO_4_ pH 7.4, 4 M guanidinium hydrochloride (GuHCl), and 5 mM imidazole) and stored at −80°C for future purification.

All FL and truncated constructs were purified by Ni-NTA chromatography followed by size exclusion chromatography. To begin, cells were resuspended in lysis buffer and sonicated on ice. The lysate was clarified by centrifugation at 15, 000 RPM for 30 minutes. The clarified lysate was run over a gravity Ni-NTA column and washed with 2 column volumes (CV) of lysis buffer and 3 CV of wash buffer (300 mM NaCl, 50 mM Na_2_PO_4_ pH 7.4, 200 mM arginine, 10% glycerol, and 5 mM imidazole). The protein was eluted with elution buffer (300 mM NaCl, 50 mM Na_2_PO_4_ pH 7.4, 200 mM arginine, 10% glycerol, and 500 mM imidazole), concentrated, and run over an S75 HiLoad 26/60 gel filtration column pre-equilibrated in 300 mM NaCl, 50 mM Na_2_PO_4_ pH 7.4, 200 mM arginine, 10% glycerol and 5 mM imidazole. The fractions containing the protein were collected, concentrated, and stored at −80 °C.

### Inducing LLPS

All FL and truncated constructs were concentrated to 50-75 μM and dialyzed into phase separation buffer (100 mM KCl, 20 mM Na_2_PO_4_ pH 7.4, and 5 mM β-mercaptoethanol (BME)). To induce phase separation, the dialyzed proteins were warmed up to their respective transition temperatures until the solution went from clear to turbid without any visible aggregates. Protein concentrations post-phase separation were taken under non-phase separating conditions such as low temperature and low concentration (i.e. when the solution is not visibly turbid).

### Performing turbidity assays using UV-Vis spectroscopy

In turbidity assays using UV-Vis spectroscopy, LLPS was examined by monitoring turbidity through light scattering at specific wavelengths over a range of temperatures (10 – 70°C). To begin, purified proteins were dialyzed in phase separation buffer for final concentrations between 5-50 μM. When protein and DNA were being studied for their ability to undergo co-LLPS, final concentrations of 5 μM and 1 μM were used, respectively (see *Oligo designs used for co-LLPS studies of MBD2 and MBD3 variants with DNA* below). Additionally, the DNA is buffer exchanged into phase separation buffer to ensure that differences in light scattering are not due to differences between buffers. To study the co-LLPS of two proteins, equimolar amounts of protein were mixed for a final concentration of 5 μM.

The cuvettes to be put into the spectrophotometer were pre-chilled at the desired starting temperature, loaded with a well-mixed solution of protein and/or DNA for a final volume of 70 μL and incubated in the spectrophotometer for 5 minutes before the start of the run. Scattering between 300-600 nm was recorded as a function of temperature by an Agilent Cary 3500 UV-Vis Spectrophotometer Multicell Peltier using a temperature ramp rate of 5°C per minute ramping from 10°C to 70°C. To ensure reproducibility and consistency, at least three replicates were run for each protein concentration. The standard deviations from the mean of the replicates are indicated by shading above and below the curve. Controls performed included buffer alone and buffer with DNA to ensure that they do not have an absorbance at scattering wavelengths. GraphPad was used to plot and analyze the data.

### Visualizing LLPS *in vitro* using DIC microscopy

Phase-separated proteins were visualized for the presence of droplets under differential interference contrast (DIC) microscopy using the Zeiss Axio Imager Z1 Upright Trinocular Fluorescence Microscope (0.0645 microns/pixel). 2 μL of protein at a desired concentration were pipetted onto a 25 x 75 x 1.0 mm glass slide (Fisherbrand, Pittsburgh, PA) and covered with a 20 mm round glass cover glass (CELLTREAT Scientific Products, Pepperell, MA). Droplets were viewed using a 100X oil immersion objective at room temperature.

### Oligo designs used for co-LLPS studies of MBD2 and MBD3 variants with DNA

The oligo designs were based on the promoter region of GSTP1. The sense and antisense strands of the oligos below were synthesized, annealed, and methylated by Integrated DNA Technologies (IDT). A final concentration of 1 μM was used for all DNA co-phase separation experiments. Model_GST_SEQ_5CG: CCCCTG**CG**ATGTCC**CG**TGGCC**CG**AGGCCT**CG**CAGCA**CG**TTGCCTG Model_GST_SEQ_2CG: CCCCTG**CG**ATGTCCCTGGGCCCAGGGCCCTGCAGCA**CG**TTGCCTG

### Expression of MBD2 and MBD3 in NIH-3T3 fibroblast cells

NIH-3T3 fibroblast cells were cultured in DMEM supplemented with 10% (v/v) fetal bovine serum (FBS), 1× 100 IU/mL penicillin and 100 μg/mL streptomycin, and 1X (2 mM) L-glutamine with 5% CO_2_ at 37°C. MBD2 and MBD3 codon-optimized for mammalian expression were cloned into a pcDNA3.1 vector tagged with eGFP or mCherry, and prepared with an endonuclease free maxi prep (Qiagen). Cells were grown to 80% confluency and transfected with 100 ng/uL of either plasmid using Lipofectamine 3000 per manufacturer’s instructions in DMEM. The transfection efficiency normally achieved for these cells is 70-99% by immunofluorescence (34). Live-cells were imaged in phenol-red free DMEM (supplemented as above) buffered with 10 mM HEPES (pH 7.4) at 37°C. Temperature was maintained with a stage heater insert (OKO labs, Ambridge, PA). 100k cells were plated in glass-bottom 35 mm dishes (MatTek, Ashland, MA). To visualize nuclei, DAPI (in 1X PBS) was added to the final concentration of 3 ug/µL to dishes 5 min before acquisition.

In addition, NIH-3T3 cells were grown as above and fixed by washing cells into 0.3% glutaraldehyde and 0.25% Triton X-100 diluted in 1X PBS and ultimately fixed in 2% glutaraldehyde for 8 min. Autofluorescence was quenched with freshly prepared 0.1%_(w/v)_ sodium borohydride. Coverslips were blocked for 1 hour in 1% BSA_(w/v)_ diluted in 1X PBST (1X PBS and 0.1% Tween-20), washed three times in 1X PBST and probed with 1X DAPI solution to stain nuclei. After 1 hour, coverslips were washed and mounted in a drop of AquaMount. Mounted slides were stored in the dark until they were imaged via laser scanning confocal microscopy (Leica SP8) equipped with a 4.2 MP CMOS camera, a 405 nm laser and an adjustable white light laser (470-670 nm) and HyD detectors using a PlanApo 100X oil objective.

### Fluorescence recovery after photobleaching (FRAP) in cells

Live NIH-3T3 cells transfected with EGFP-MBD2 or EGFP-MBD3 protein were visualized for protein droplets at 100X using a Leica SP8 confocal microscope. Protein condensates were completely bleached using a 488 nm laser at 50% laser power. Recovery of GFP signal to the bleached area was monitored every second for up to 60 seconds after bleaching. Image montages showing the recovery of a representative FRAP experiment were generated using FIJI/ImageJ (35).

### Coarse-grained molecular dynamics simulations and analysis

The coarse-grained (CG) coexistence simulations of MBD2 and MBD3 with and without DNA were performed in the HOOMD-Blue 2.9.7 software package using the slab geometry as described in previous studies (36–39). The MBDs of MBD2 and MBD3 proteins were constructed using the available structural models of these domains (PDB ID: 6CNP and 2MB7) (22, 40). Disordered regions were connected to the MBDs using MODELLER (41). The folded domains of MBD2 and MBD3 were constrained using the hoomd.md.constrain.rigid function (42, 43) while disordered regions remained flexible, following our previously established approach for multidomain proteins (44, 45). The proteins and DNA were modeled using the one-bead-per-residue HPS-Urry model and the recently developed two-bead-per-nucleotide DNA model, respectively (37, 46). The CG coexistence simulations were initiated in box dimensions of 180×180×1260 Å^3^. The number of chains of MBD2 and MBD3 (50 and 66, respectively) were chosen to give similar protein concentrations. The simulation box size and the number of protein chains were selected to account for the potential impact of finite-size effects as done in our previous work (39).

In all CG simulations, we use a runtime of 5 μs in an NVT ensemble using a Langevin thermostat with a friction factor γ = m_AA_/τ. Here, m_AA_ and τ are the mass of each amino acid bead and the damping factor (set to 1000 ps), respectively. The time step was set to 10 fs. As the coexistence density in the dilute phase was too low at 300 K, we simulated all systems at 320 K to compare phase separation between the two FL MBD proteins and their respective truncations similar to our approach for HP1α (38). All the CG simulations were conducted at 100 mM salt concentration. When calculating the density profiles and contact maps, the first 1 μs of the trajectory was excluded and treated as an equilibration period. In the contact analyses, two residues were considered to be in contact if the distance between them was less than 1.5 of the arithmetic mean of their Van der Waals (vdW) radii. Snapshots of the simulations were visualized using VMD (47).

## RESULTS AND DISCUSSION

### MBD2 and MBD3 form dynamic condensates *in cellulo*

Previous *in cellulo* experiments have demonstrated that MeCP2 can form dynamic condensates (5, 9, 19). MBD2 and MBD3 LLPS-like condensates have been previously recognized as puncta *in cellulo* when LLPS was not widely recognized to occur in a biological context (48, 49). However, the study of MBD2 and MBD3 LLPS has not been further explored since those initial observations. Disordered, multivalent molecules that facilitate transient homo- and heterotypic interactions are likely to phase separate. Because MBD2 and MBD3 display a high degree of disorder and contain multiple modular domains (Figures 1A-C) known to interact with DNA and protein subunits of repression complexes, similar to MeCP2, these proteins are likely to undergo LLPS (48, 50). Additionally, MBD2 and MBD3 contain post-translational modification sites (PTMs) along their disordered regions that, when modified, can change their interactive properties and induce conformational changes that could influence condensate structure and dynamics (26, 50–53). Therefore, we wanted to confirm the observance of this phenomenon *in cellulo* by recreating what others have previously characterized as puncta. EGFP-tagged MBD2 and MBD3 were overexpressed in NIH-3T3 fibroblast cells and visualized under confocal fluorescence microscopy (Figure 1D). We observed the presence of MBD2 and MBD3 spherical puncta (Figure 1D). To ensure these are phase-separated, dynamic condensates, we performed FRAP experiments and demonstrated that these MBD2 and MBD3 puncta are indeed dynamic protein condensates with liquid-like properties (Figure S1). Overall, these results show that MBD2 and MBD3 are both dynamic components of heterochromatin condensates in live NIH-3T3 cells similar to what was observed for MeCP2 (5, 9).

### Inducing MBD2 and MBD3 LLPS *in vitro* and *in silico*

To further confirm the condensates observed *in cellulo* are mediated by LLPS and explore the homotypic and heterotypic interactions that drive the formation of these droplets, we performed a combination of *in vitro* experiments and *in silico* simulations. The interactions that drive MBD2 and MBD3 LLPS may be mediated differently due to differences in their amino acid sequences and domain architecture. A more in-depth analysis of their amino acid composition can help identify sequence motifs or features important in determining the interactions driving LLPS that ultimately influence their own specific roles as transcriptional regulators. Not only are we looking for features that are unique to each protein, but additionally, we can also uncover features that might be conserved within the MBD family of proteins.

As *in cellulo*, MeCP2 can phase separate *in vitro* with droplets exhibiting LLPS behavior such as flowing and fusing as well as sensitivity to differing protein and salt concentrations. MeCP2’s electrostatically-driven LLPS is attributed to its high proportion of charged amino acids (33%) (5, 19). Similarly, MBD2 and MBD3, enriched in Arg, Lys, Glu, and Asp, contain 26-32% charged residues. These three MBD proteins also have a comparable composition of other amino acid types, including aromatic (4-7%), aliphatic (22-28%), polar (19-22%), and other residues (12-24%) (Figures 1C and 1E). Given the high fraction of charged residues in MBD2 and MBD3, it is plausible that multivalent, electrostatic interactions primarily drive their LLPS behavior. However, these interactions might be modulated differently in each protein due to variances in their amino acid sequences and how their respective folded domains mediate intra- and interdomain interactions.

We wanted to experimentally validate our *in cellulo* results and sequence analyses by inducing the phase separation of FL MBD2 and MBD3 *in vitro*. The extent of disorder of these proteins interspersed with regions of folded or partially folded domains, however, has made it challenging to recombinantly express and purify at the high concentrations required for biophysical characterization without any fusion tags and crowding agents to avoid artifacts. Typically, studies on intrinsically disordered proteins that undergo phase separation involve small, truncated regions of proteins that are more amenable to biophysical studies. Previous studies of MBD2 and MBD3 have characterized truncated forms of MBD2 and MBD3, but still suffered from low yields due to degradation and aggregation (30, 31, 56). We have also encountered extensive protein degradation but have adapted a protocol effective for intrinsically disordered proteins and optimized it for MBD2 and MBD3 (32, 57). As a result, we have obtained large amounts of protein suitable for phase separation studies (See *Expression and purification of MBD2 and MBD3 and their respective truncations* under Materials & Methods).

Upon exchanging purified MBD2 and MBD3 into a phase separation buffer, we observed the appearance of a turbid solution that contained microscopic droplets (Figure 2A). As expected, the droplets scaled in size and number with increasing protein concentration. We also performed turbidity assays as a function of temperature as it is a well-studied factor that regulates protein phase behavior (58–60). Depending on the types of interactions driving LLPS, proteins will demix in response to changes in temperature. An increase or decrease in temperature gives rise to phase transitions with either lower critical solution temperature (LCST) or upper critical solution temperature (UCST), respectively (61). More specifically, UCST behavior is thought to result from both entropic and enthalpic contributions stemming from electrostatic and hydrophobic interactions, as well as cation-π and π-π stacking interactions, whereas LCST behavior stems mainly from the gain in solvent entropy (58). Systems can also exhibit a dual-phase behavior where both UCST and LCST behaviors are observed within the accessible temperature range in experiments (58). Our turbidity assays as a function of temperature revealed that both MBD2 and MBD3 exhibit a dual LCST/UCST phase behavior in a concentration-dependent manner (Figure 2B and C). As temperature and protein concentration increased, phase separation correspondingly increased as indicated by the growing amplitudes of the phase transition curves and earlier onset transition temperatures (Figures 2B and C). However, it is important to note that MBD2’s biphasic phase behavior is more pronounced than MBD3’s as it peaks earlier at approximately 55°C. This may be due to the more distinct block-copolymeric architecture with its additional GR-rich N-terminal domain compared to MBD3. Furthermore, these dual transitions indicate that multiple interaction types, including solvent effects, might be at play within certain temperature ranges.

**Figure 2.**
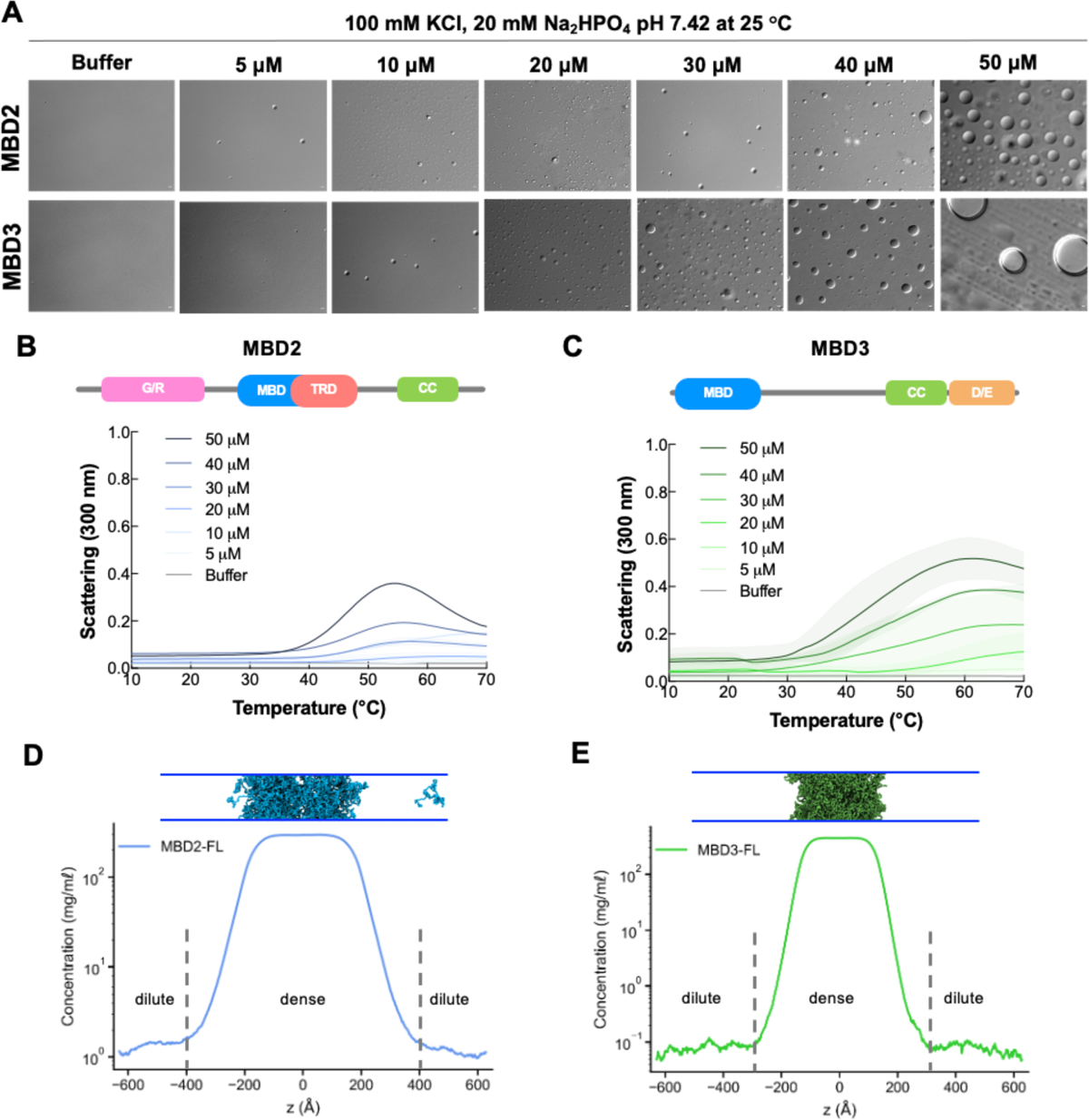
MBD2 and MBD3 undergo LLPS *in vitro* and *in silico*. **A.** DIC microscopy images of MBD2 and MBD3 (5-50 μM) in phase separation buffer (100 mM KCl, 20 mM Na2HPO4 pH 7.4) at 25 °C. Images were taken under 100x magnification. Scale bar = 2 µm. **B and C**. UV-Vis absorption spectra of MBD2 and MBD3 in phase separation buffer as a function of temperature at increasing concentrations (5-50 μM). Above the spectra is a schematic of each protein’s domain architecture. The shading around each curve represents the standard deviation from the mean absorbance from technical replicates. Scattering, or the turbidity of the solution, is used as a proxy for LLPS. **D and E**. Density profiles and condensate snapshots in the CG co-existence simulations of MBD2 and MBD3. The CG coexistence simulations were conducted using the HPS-Urry model at 320K

To gain insights into the molecular interactions driving the phase separation of MBD2 and MBD3, we conducted CG phase coexistence simulations using the HPS-Urry model with the slab geometry (see Methods). Both FL MBD2 and FL MBD3 were modeled using MODELLER (41). The folded MBD was kept rigid to prevent protein unfolding by applying a rigid body constraint while the rest of the chain remained flexible (see Materials and Methods). We simulated the systems at 320 K and plotted the protein densities as a function of the z-coordinate, effectively distinguishing between dense and dilute phases (Figures 2D, E, Movie S1). Our results show that both MBD2 and MBD3 formed stable condensates, indicated by the flat density profiles and snapshots in Figures 2D and 2E. To elucidate the interactions facilitating the phase separation of MBD2 and MBD3, we generated one- and two-dimensional intermolecular contact maps based on van der Waals (vdW) contacts within the condensed phase (Figures S2 A, B). These maps highlight high contact-prone regions along the protein sequence. Notably, the acidic tail in MBD2 strongly interacts with its GR-rich region (Figure S2 A) while in MBD3, the acidic DE tail shows a marked preference for interactions with both the positively charged MBD and positively charged N-terminal region of its IDR (Figure S2 B). In both cases, these can be attributed to electrostatic attractions between positively charged K/R and negatively charged D/E residues.

### Defining the homotypic interactions driving MBD2 and MBD3 LLPS

How do the structural elements within MBD2 and MBD3 contribute to their ability to phase separate? Like the interplay observed between the IDRs and folded domains within HP1α and MeCP2 that give rise to their LLPS behavior (5, 17, 39), we similarly aim to understand how the interactions between the folded and disordered regions within MBD2 and MBD3 contribute to their LLPS. We designed MBD2 and MBD3 truncations that include or exclude important sequence-based features or structural and functional regions to test their influence on LLPS propensity (Figures 3A and Figure 4A).

**Figure 3.**
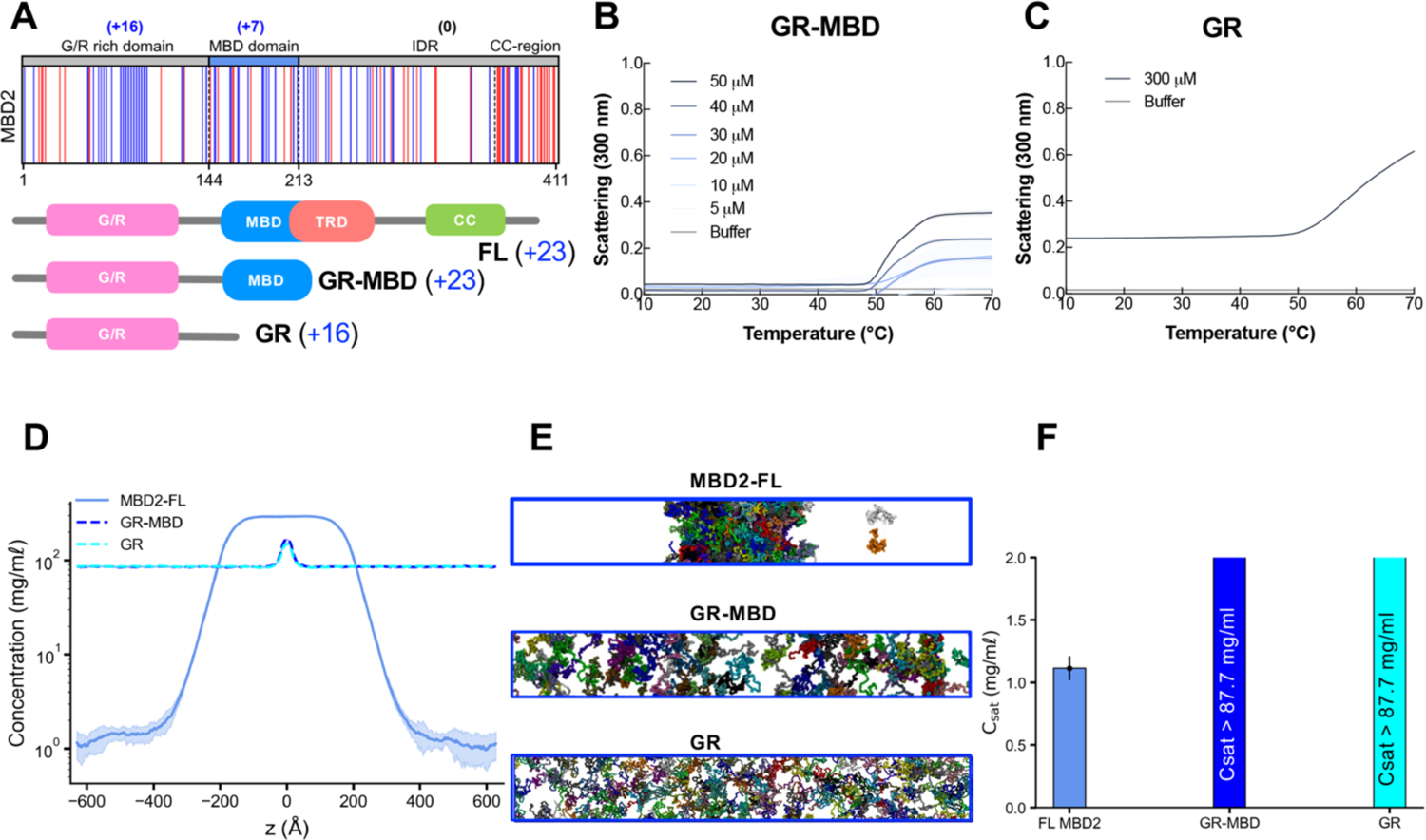
Domain contribution in MBD2 LLPS. **A.** Charge distribution in the sequence (blue = positively charged residues, red = negatively charged residues) and domain architecture of MBD2 truncations. **B and C**. UV-Vis absorption spectra of truncations of MBD2, GR-MBD, and GR in phase separation buffer as a function of temperature at increasing concentrations. GR did not have a noticeable absorbance between 5-50 μM but was shown to phase separate around 300 μM under these conditions. The shading around each curve represents the standard deviation from the mean absorbance from technical replicates. **D, E, and F**. Density profiles, representative snapshots (the system was colored by protein chains), and the saturation concentration in the CG coexistence simulations of MBD2 and its truncations. The error bars represent the standard deviation from triplicate simulation sets. The CG coexistence simulations were conducted using the HPS-Urry model at 320K.

**Figure 4.**
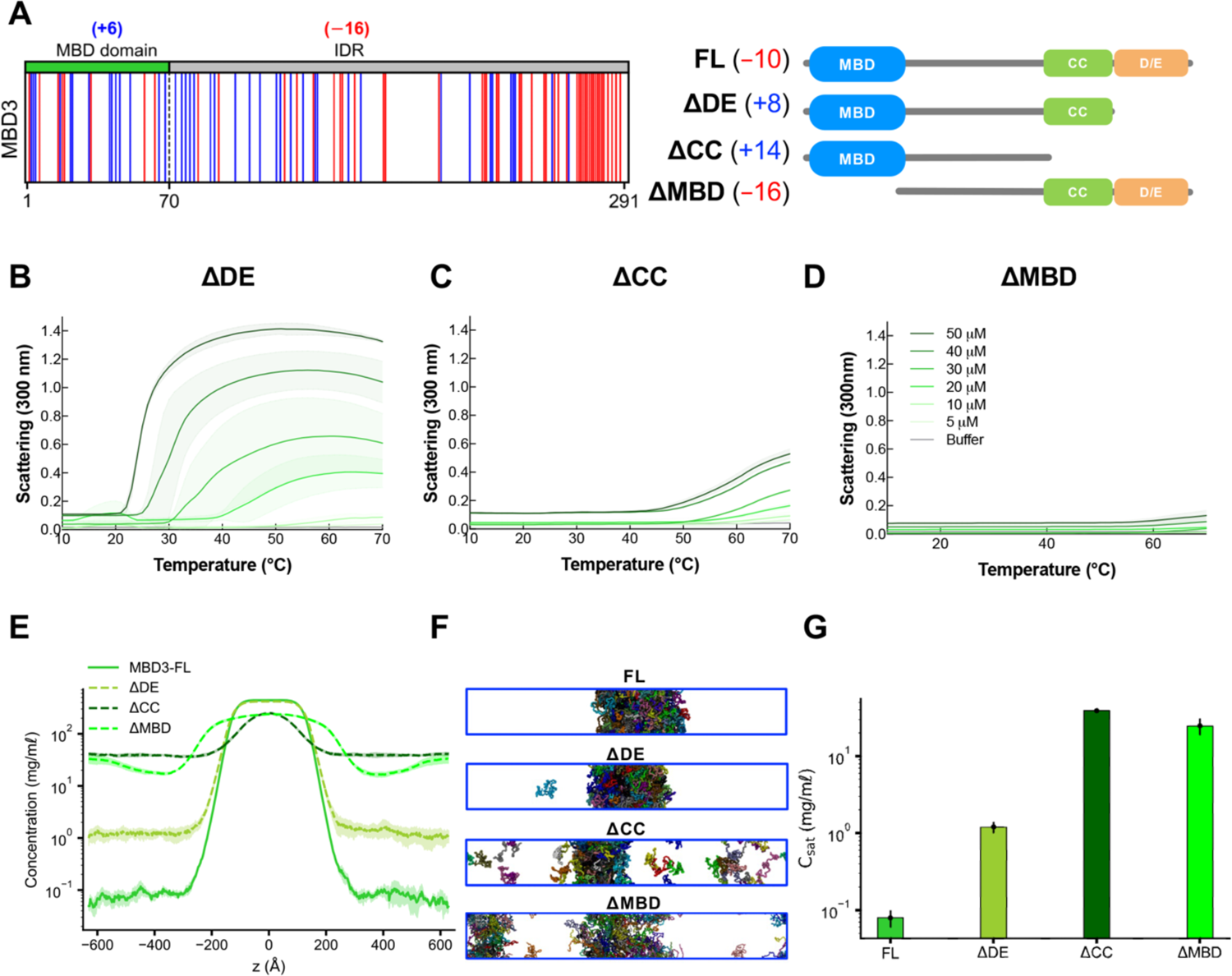
Domain contribution in MBD3 LLPS. **A.** Charge distribution in the sequence (blue = positively charged residues, red = negatively charged residues) and domain architecture of MBD3 truncations. **B, C, and D**. UV-Vis absorption spectra of truncations of MBD3, which include ΔMBD, ΔDE, and ΔCC, in phase separation buffer as a function of temperature at increasing concentrations (5-50 μM). The shading around each curve represents the standard deviation from the mean absorbance from technical replicates. **E, F, and G**. Density profiles, representative snapshots (the system was colored by protein chains), and the saturation concentration in the CG coexistence simulations of MBD3 and its truncations. The error bars represent the standard deviation from triplicate simulation sets. The CG coexistence simulations were conducted using the HPS-Urry model at 320K.

MBD2 phase separation could potentially be mediated by interactions from the MBD, coiled-coil domain, and/or charge-enriched disordered regions. Turbidity assays show that the removal of the coiled-coiled domain and large IDR in MBD2’s C-terminus diminishes LLPS propensity as the approximate temperature of phase separation onset becomes higher than that of the FL MBD2 (Figures 2B and 3B). The amplitudes of the curves remain roughly the same, but the biphasic nature observed in FL MBD2 is absent, suggesting a synergistic interplay between MBD and IDR domains in facilitating the phase separation of MBD2 (Figures 2B and 3B). When the MBD is additionally removed, phase separation is completely diminished within the concentration range of 5 μM to 50 μM, indicating that the MBD is also important in driving LLPS (Figure 3C). A similar effect was observed for MeCP2 (5). Furthermore, we performed the CG coexistence phase simulations of the MBD2 truncations, maintaining similar initial protein concentrations to those of the FL MBD2. We also calculated the coexistence density in the dilute phase (referred to as saturation concentration, C_sat_) to assess the phase separation propensity of FL MBD2 and its truncations. We found that neither the G/R-rich domain alone nor in conjunction with the MBD underwent phase separation (Figures 3D-F). These observations are consistent with the experimental data and underscore the critical intermolecular interactions between the MBD, coiled-coil domain, and the GR-rich region in driving MBD2 phase separation (Figures 3A and S2 A).

MBD3 exhibits a charge patterning in its MBD and C-terminus that is distinct from MBD2. Negatively charged residues (D/E) cluster along the C-terminal end of MBD3 that contrast with the dispersed positively charged residues (K/R) on the MBD and the N-terminal region of the IDR (Figure 4A). This charge distribution, as suggested by our contact analysis (Figure S2 B), appears critical for MBD3’s ability to phase separate. Based on these observed features and important structural and functional regions within MBD3, we explored the specific contributions of the MBD, coiled-coil domain, and D/E-rich tail to MBD3’s phase behavior. We created truncations that exclude the D/E-rich region (ι1DE), coiled-coil domain (ι1CC), and the MBD (ι1MBD), and tested their LLPS behavior (Figure 4A).

Based on the turbidity assays in Figure 4, when the D/E-rich region in MBD3’s C-terminal domain is eliminated, LLPS is enhanced, as seen in the enlargement of amplitudes in the phase transition curves and lower transition temperatures compared to the FL protein (Figures 2B and 4B). When the CC domain, a significant portion of C-terminal IDR, is eliminated, LLPS is reduced, as evidenced by the higher transition temperatures and smaller amplitudes, suggesting the importance of the coiled-coil domain in the LLPS of MBD3 (Figure 4C). Furthermore, under our tested conditions, phase separation is completely diminished when the MBD is removed, while leaving the C-terminal IDR intact, indicating the importance of the interactions between the MBD and the C-terminal IDR (Figure 4D). To further investigate the contributions of specific regions to MBD3’s propensity for phase separation, we performed CG coexistence phase simulations of the truncations at 320K. The deletion of the D/E-rich tail shifted the net charge from negative (q=-10) to positive (q=+8), yet the condensate remained stable. This change led to a shift in dominant interactions from negatively charged residues (e.g., Glu and Asp) to predominantly hydrophobic and positively charged residues (e.g., Leu and Lys), likely due to their prevalence in the sequence (Figures S2 C-E). However, our computational model does not fully account for temperature-dependent interactions, which appear critical in promoting phase separation of the ΔDE truncation, as observed experimentally. Furthermore, the subsequent removal of the coiled-coil domain resulted in an increase in the C_sat_ by more than two orders of magnitude. Similarly, deleting the MBD increased Csat significantly compared to FL MBD3. Consistent with experimental results, these findings underscore the pivotal roles of the MBD and coiled-coil domains in MBD3’s phase separation. Additionally, hydrophobic residues, constituting about 30% of the MBD3 sequence, may also contribute to its LLPS, akin to the role of methionine residues in the LLPS of TDP-43 (62).

Thus far, we have analyzed the homotypic interactions that drive the LLPS of MBD2 and MBD3 individually. Our combined computational and experimental analyses reveal the influence of electrostatic and hydrophobic interactions in MBD2 and MBD3 LLPS. Specifically, the domain contributions in each protein, including the MBD and various charged regions, are essential in dictating their respective LLPS behaviors.

### Inducing MBD2 and MBD3 co-phase separation and exploring their co-localization within condensates

MBD2 and MBD3 coordinate with each other to perform their transcriptional repression duties (15, 27, 28). However, the potential of these proteins to co-localize and undergo co-phase separation, a phenomenon critical for various cellular processes remains unexplored (63–65). The contrasting charge profiles of MBD2 (q=+23) and MBD3 (q=-10) suggest a potential for electrostatic complementarity. Based on these observations, MBD2 and MBD3 may have the ability to co-localize and co-phase separate.

To test this, we first performed CG coexistence simulations of MBD2 and MBD3 mixed at an equimolar ratio. We found that the MBD2/MBD3 mixture readily formed a stable condensate (Figure 5A, Movie S2). Both MBD2 and MBD3 have a strong homotypic affinity and were able to phase separate on their own and cooperatively form heterotypic condensates. The acidic tail of MBD3 shows preferential cross-interactions with the G/R-rich region and positively charged clusters distributed in the MBD of MBD2 (Figure 5B). We next varied the concentration ratios of MBD2 to MBD3 in the mixture while maintaining a constant total protein concentration in the CG coexistence simulations and constructed the phase diagram of this two-component system to investigate the co-phase separation of MBD2 and MBD3 (Figure 5C). We observed that the concentration of MBD3 in the dilute phase remained consistently low across all tested ratios whereas the dilute concentration of MBD2 exhibited a significant increase starting from the ratio of 2:1 (lower left corner of the phase diagram). This rise suggests a nuanced interplay between MBD2 and MBD3, potentially altering phase separation behavior, presumably due to the unbalanced net charge resulting from the increased proportion of MBD2 (+23) relative to MBD3 (−10). Notably, at the highest ratio of MBD2 to MBD3, the dilute concentration of MBD2 decreased to approximately half of its concentration when present alone (Figures 3F, 5C). This suggests that even a minimal presence of MBD3 can substantially stabilize the phase separation of MBD2.

**Figure 5.**
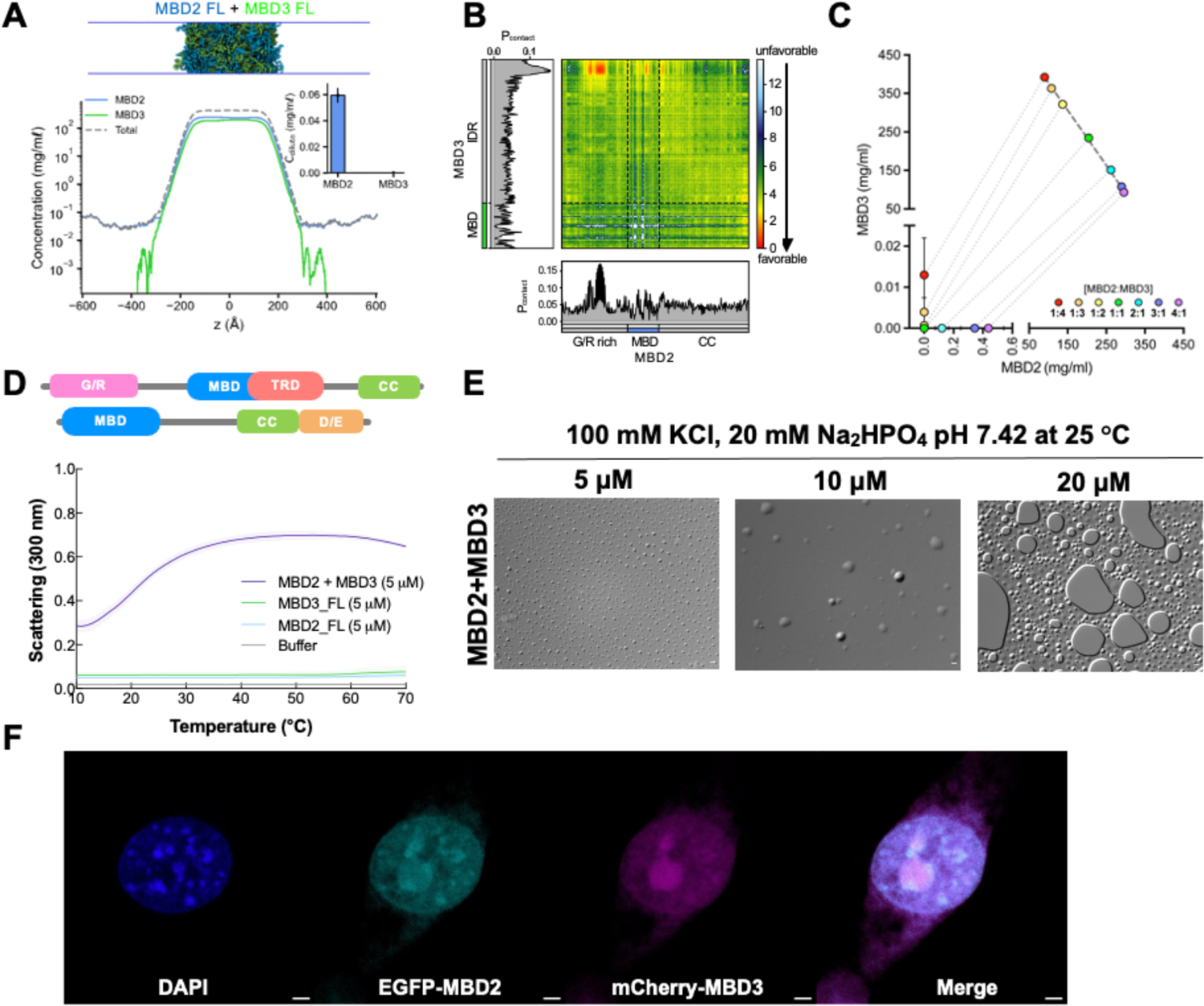
Heterotypic interactions promote MBD2 and MBD3 co-localization. **A.** Density profile, a representative snapshot of MBD2-MBD3 condensate, and the dilute concentrations of MBD2 and MBD3 (inset) in the equimolar CG coexistence simulation. **B.** Intermolecular contacts between MBD2 and MBD3 within the condensate along the sequence of each protein. The CG coexistence simulations were conducted using the HPS-Urry model at 320K. **C**. Multicomponent phase diagram of MBD2 and MBD3 co-phase separation. The concentration ratios of MBD2 to MBD3 used to generate the phase diagram are color-coded in the legend. The tie line (gray dotted line) connects the concentrations in the dense and dilute phases for each ratio. Phase existence line (black dashed line) defines the high-concentration arm of the phase diagram. The error bars represent the standard deviation from triplicate simulation sets. The CG coexistence simulations were conducted using the HPS-Urry model at 320K. **D.** UV-Vis absorption spectra of FL MBD2 and FL MBD3 individually (5 μM) and mixed at an equimolar ratio (final concentration of 5 μM) in phase separation buffer as a function of temperature. **E.** DIC microscopy images of FL MBD2 and MBD3 mixed at equimolar ratios (final concentrations of 5, 10, or 20 μM) in phase separation buffer (100 mM KCl, 20 mM Na_2_HPO_4_ pH 7.4) at 25°C. Images were taken under 100x magnification. Scale bar = 2 µm. **F**. Live 3T3 fibroblast cells co-transfected with eGFP-tagged FL MBD2 and mCherry-tagged FL MBD3 viewed under different channels to visualize DNA with DAPI (blue), MBD2 (cyan), and MBD3 (magenta) and then a merged view. Scale bar = 2 µm.

We first assessed the ability of FL MBD2 to undergo co-phase separation *in vitro* with FL MBD3, thereby providing additional insights into the heterotypic interactions observed in our *in silico* findings. Turbidity assays, conducted as a function of temperature, demonstrated that the MBD2 and MBD3 complex exhibits a dual LCST/UCST phase behavior, which is dependent on both temperature and concentration, similar to the behavior of individual proteins. As the temperature and protein concentration increased, the phase separation also enhanced (Figures 5D and E). This trend was evidenced by lower transition temperatures and increasing amplitudes in the phase transition curves, with light scattering measurement starting at approximately 0.3 at 0°C and increasing to about 0.6 (Figure 5D). At a concentration of 5 μM, both MBD2 and MBD3 individually demonstrated negligible turbidity. However, their mixture resulted in a significant increase in LLPS, suggesting a synergistic effect rather than a merely additive response. Consistent with the turbidity assays, the mixing of equimolar amounts of phase-separated MBD2 and MBD3, at final concentrations of 5, 10, and 20 μM, lead to a turbid solution containing microscopic where both size and number increased with the protein concentration. The mixing of these two proteins significantly enhances LLPS, as evidenced by the larger droplets observed at 20 μM compared to those formed by the individual proteins alone (Figures 2A and 5F). In alignment with our *in vitro* findings, MBD2 and MBD3 not only co-localize but also co-phase separate *in cellulo* (Figure 5F).

The oligomeric states of these proteins, as well as the relative affinities between segments and their flexibility in interactions, remain undetermined. According to our *in silico* simulations for the individual proteins and their co-localization, electrostatic and hydrophobic interactions, whether from IDRs and/or within folded domains, are the primary drivers of the co-LLPS of MBD2 and MBD3. In our turbidity assays involving the mixing of MBD2 and MBD3, the removal of MBD3’s acidic tail (ΔDE) resulted in a complete loss of LLPS (Figure S3A). This observation suggests that MBD3’s acidic tail is important in mediating the LLPS of the complex, likely through interactions with the MBD2’s G/R-rich domain, as shown in our *in silico* predictions (Figure 5B). When the coiled-coil domain was additionally removed (ΔCC), the complex remained unable to undergo LLPS (Figure S3B). Notably, the complex exhibiting a phase-separation profile most similar to that of the two FL proteins is the FL MBD2 combined with MBD3-ΔMBD. Although this mixture did not initiate phase separation as immediately as the two FL proteins, its LLPS began at approximately 20°C and achieved the highest scattering intensity of ∼0.8 (Figure S3C). The results suggest that the acidic tail of MBD3 is essential for facilitating the co-LLPS of MBD2 and MBD3.

We further explored the ability of FL MBD3 to co-phase separate with truncated forms of MBD2, specifically the G/R-rich domain (MBD2-GR) and a construct containing both the G/R-rich domain and the MBD (MBD2-GR-MBD). Turbidity assays indicated that removing the MBD and C-terminal domain of MBD2 significantly impairs its ability to co-phase separate with FL MBD3, compared to assays with both proteins in their FL forms. Specifically, co-phase separation occurred only at elevated temperatures (∼30°C) and reached a lower maximum scattering intensity (∼0.4), as shown in Figure S3D. These findings corroborate previous observations, suggesting that the acidic tail of MBD3 and the G/R-rich region of MBD2 are crucial for co-phase separation. We also observed that including the MBD domain in the MBD2-GR construct (MBD2-GR-MBD) further reduced co-phase separation compared to FL MBD3 and MBD2-GR alone (Figure S3E). Remarkably, when the IDR of MBD3 (MBD3-ΔMBD) was mixed with MBD2’s G/R domain, no LLPS was observed (Figure S3F), underscoring that while the presence of oppositely charged regions is necessary, at least one intact MBD domain is essential for robust co-phase separation. Our findings reveal the cooperative roles of disordered and folded domains in mediating MBD2 and MBD3 co-phase separation, demonstrating an intricate balance of structural elements crucial for their functions.

### Investigating DNA’s influence on MBD2 and MBD3 LLPS and co-localization

In addition to the MBD that binds both methylated and unmethylated CpG-containing DNA, albeit with lower affinity for unmethylated CpG, the MBD protein family is known to interact with DNA using DNA binding motifs embedded in their disordered regions (21, 27, 66–69). It has been demonstrated that the addition of DNA increases the propensity for MeCP2 to phase separate with a concomitant increase in droplet size, depending on the length and methylation status of the DNA substrates (5, 19). MBD2 binds to double-stranded, methylated DNA *in vivo* and *in vitro* (27, 66–69). Therefore, we wanted to know whether the addition of DNA would enhance its ability to undergo LLPS. Additionally, we wanted to determine whether there was a difference in LLPS propensity between methylated and unmethylated DNA and the number of methyl groups. We expect, as suggested by its biological preferences, that MBD2 LLPS will be most enhanced by methylated DNA containing more methylated sites. We have designed DNA oligos containing either 2 or 5 CpG sites based on the CpGs island sequence near the *GSTP1* promoter known to be regulated by MBD2 (See *Materials & Methods*) (70). Based on the turbidity assays, the addition of DNA, whether unmethylated or methylated at 2 or 5 CpG sites, enhanced MBD2 LLPS (Figure 6A). It should be noted that we performed the turbidity assay with a low concentration (5 μM) of MBD2. The LLPS of MBD2 at 5 μM without DNA is almost negligible. The addition of DNA greatly enhances LLPS to an observed initial scattering between 0.4 and 0.6, while MBD2 alone, at the same concentration, has an initial scattering near 0. Moreover, MBD2 with DNA reaches a higher turbidity than MBD2 alone at its highest concentration (Figure 6A and 2B). The presence of DNA promotes multivalency and interacts cooperatively with MBD2, leading to robust phase separation. This LLPS is likely facilitated by interactions between DNA and the positively charged N-terminal region, MBD, or K/R clusters with the C-terminus of MBD2. However, the GR-rich region alone with DNA was not sufficient to induce LLPS (Figure S4A). It is important to note that our observations may not reveal significant differences between various DNA substrates, potentially due to the limited sensitivity of the assay or because the level of DNA methylated sites is not sufficient to induce a robust change in LLPS propensity.

**Figure 6.**
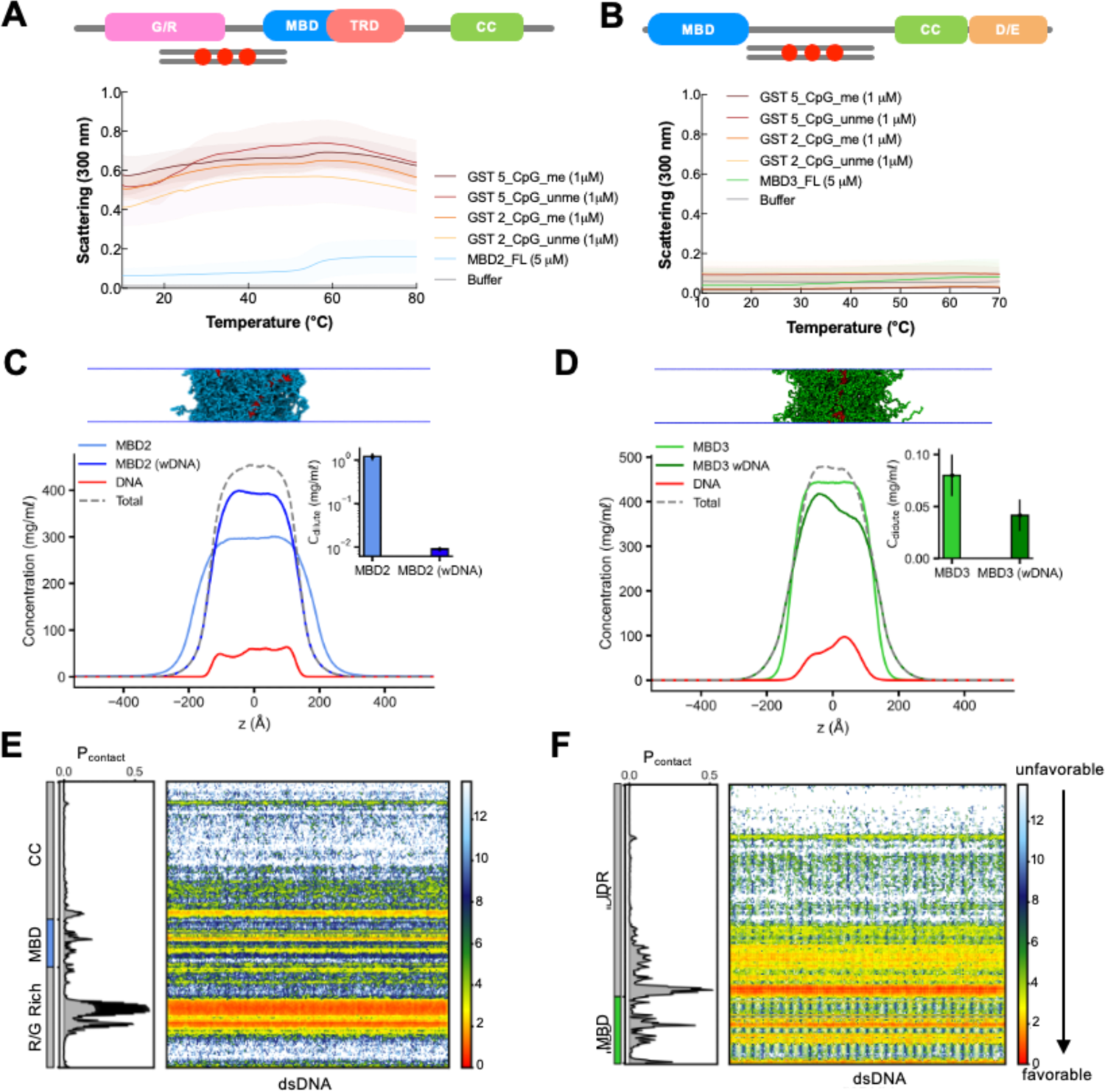
MBD2 and MBD3 LLPS with DNA. **A and B.** UV-Vis absorption spectra of FL MBD2 (left) and FL MBD3 (right) individually and mixed with either unmethylated or methylated DNA that contain either 2 or 5 CpG sites in phase separation buffer as a function of temperature. Protein and DNA concentrations remained constant at 5 μM and 1 μM, respectively. The shading around each curve represents the standard deviation from the mean absorbance from technical replicates. **C and D**. Density profiles, representative snapshots of the condensate, and the dilute concentrations of MBD2 and MBD3 (inset) in the presence of dsDNA in CG coexistence simulations. The error bars represent the standard deviation from triplicate simulation sets. **E and F.** Intermolecular contacts between MBD2/MBD3 and DNA within the condensed phase. Preferential interactions are shown in red. The CG coexistence simulations were conducted using the HPS-Urry model at 320K.

While prior studies have shown that the MBD of MBD3 does not directly bind methylated DNA (67, 69), we explored whether DNA, independent of MBD binding, could enhance MBD3 LLPS using the same substrates as in MBD2’s turbidity assays. Intriguingly, our results indicate that neither methylated nor unmethylated enhanced the LLPS of MBD3 (Figure S4B), despite the 70% identity between C-terminal IDRs of MBD2 and MBD3, including predicted DNA-binding residues. However, a key difference lies in their net charge: MBD3’s C-terminus carries a strong negative charge (−16) due to its pronounced acidic tail, whereas MBD2’s C-terminus is neutral. We hypothesized that removing this acidic tail (MBD3-ΔDE) might facilitate DNA interaction and thereby enhance LLPS. Indeed, the addition of DNA, whether methylated or unmethylated, successfully induced LLPS in MBD3-ΔDE, similar to MBD2 (Figure S4B). These findings suggest that the acidic tail of MBD3 might act as a repulsive control element for DNA, preventing LLPS in FL MBD3.

Motivated by the *in vitro* results, we sought to explore the origins of the molecular interactions influencing MBD2 and MBD3 phase separation in the presence of DNA. We first employed HybridDBRpred (71), a recently improved sequence-based prediction program for DNA-binding proteins that uses annotations from both structured and disordered regions of proteins. Apart from their C-terminal D/E-rich acidic tail, both MBD2 and MBD3 contain multiple regions throughout their structure that are predicted to possess DNA binding residues (Figure S5). To test this further, we employed a recently developed nucleic acid model (46) to conduct CG coexistence simulations with the mole fraction of MBD proteins and dsDNA reflecting a 5:1 protein-to-DNA ratio akin to our experimental setup. We found that dsDNA partitioned into and stabilized the condensates of MBD2 and MBD3 (Figures 6C, D, Movie S3). Interestingly, the C_sat_ of MBD2 decreased significantly by approximately two orders of magnitude in the presence of dsDNA compared to MBD2 alone. Conversely, MBD3 showed only a slight reduction in C_sat_ within error margins, likely due to the electrostatic repulsion from its acidic tail. To further elucidate protein-DNA interactions, we computed the vdW-based intermolecular contact maps formed between MBD proteins and dsDNA as a function of residue number. In MBD2, the electrostatic interactions predominantly occurred between dsDNA and the G/R-rich region. Additionally, dsDNA showed an affinity for positively charged areas on the MBD surface and C-terminal IDR near the MBD. In contrast, these interactions are more prominent in the MBD and extended stretches of positively charged residues in the N-terminal region of the IDR of MBD3. However, the phase separation of MBD3 was less influenced by dsDNA, likely due to electrostatic repulsion from its acidic tail. It should be noted that the coexistence simulations in the presence of dsDNA were conducted at high protein concentrations, conditions under which the proteins readily undergo phase separation. Consequently, the influence of dsDNA on the phase separation of MBD3 appears to be less pronounced, as evidenced by experimental observations. Overall, our *in vitro* and *in silico* findings highlight the role of DNA in modulating the phase separation of MBD proteins, offering deeper insights into the mechanisms driving heterochromatin organization.

Building upon our findings on the co-LLPS of MBD2 and MBD3, we delved deeper into how DNA influences their co-localization. Central to our investigation were two key questions: first, the degree to which the presence of DNA affects the LLPS dynamics of this complex, and second, the variations in LLPS propensity in response to methylated versus unmethylated DNA with a particular focus on the influence of the number of methyl groups. Based on the turbidity assays, the addition of DNA, whether unmethylated or methylated at 2 or 5 CpG sites, further enhanced the LLPS of the complex (Figure 7A). Here, we achieve the greatest propensity to phase separate than the individual proteins alone at the highest concentrations used in this study, the individual proteins with DNA, or mixing of the full-length proteins. Although this may be simply due to an increase in multivalency, it may also have biological significance as all components at low concentrations enhance LLPS substantially. Our CG coexistence simulations also show that an equimolar mixture of MBD2 and MBD3 formed a stable condensate with the addition of dsDNA at a protein-DNA ratio of 5:1 (Figure 7B, Movie S4). Initially, in the absence of DNA, MBD3 preferentially localized within the condensate of the MBD2/MBD3 mixture. However, upon the introduction of DNA, a notable reversal was observed: MBD2 predominantly localized within the condensate (insets of Figures 5A, 7B), likely facilitated by its enhanced affinity for both DNA and MBD3 (Figures 7C-E). These observations underscore the role of DNA in modulating the co-localization of MBD proteins, which carries significant implications for understanding their biological functions in the context of chromatin (39, 72).

**Figure 7.**
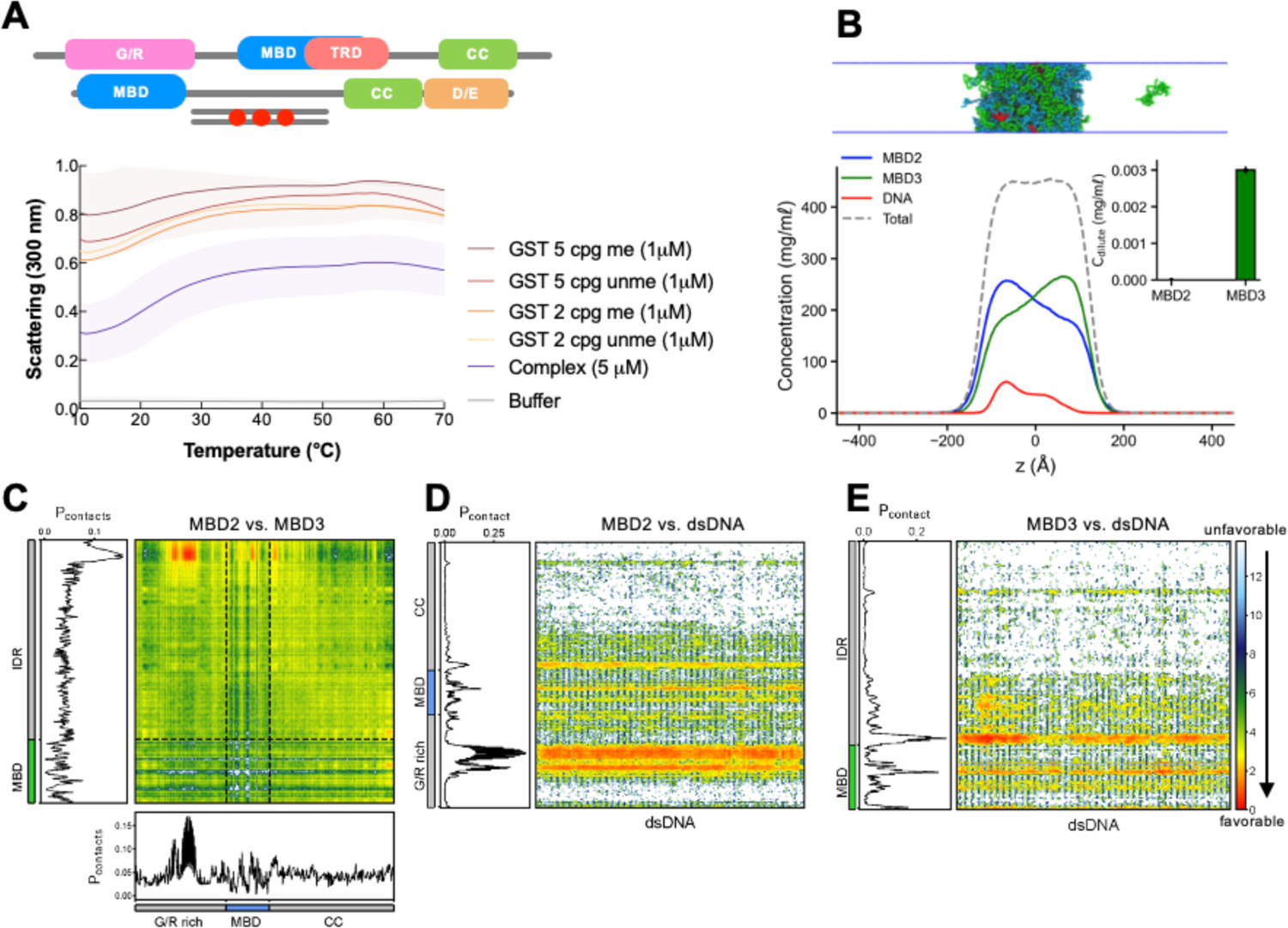
DNA’s influence on MBD2 and MBD3 co-localization. **A**. UV-Vis absorption spectra of full-length MBD2 mixed with an equimolar ratio of full-length MBD2 with either unmethylated or methylated DNA that contains 2 or 5 CpG sites. Protein concentration and DNA remained constant at 5 μM (final) and 1 μM, respectively. The shading around each curve represents the standard deviation from the mean absorbance from technical replicates. **B**. Density profiles, representative snapshots of the condensate, and the dilute concentrations of MBD2 and MBD3 (inset) at equimolar in the presence of dsDNA in CG coexistence simulations. The error bars represent the standard deviation from triplicate simulation sets. **C-E**. Intermolecular contact maps of protein-protein (MBD2 vs. MBD3) and protein-DNA (MBD2/MBD3 vs. DNA) within the condensed phase. Preferential interactions are shown in red. The CG coexistence simulations were conducted using the HPS-Urry model at 320K.

## Summary and Conclusion

Heterochromatin, traditionally viewed as a static structure characterized by its compactness and transcriptional repression, has been shown to regulate its functions spatiotemporally through the formation of condensates, displaying dynamic behavior across various lengths and timescales (8, 73, 74). Proteins associated with heterochromatin, particularly those containing IDRs such as HP1α and MeCP2, are believed to facilitate the LLPS of chromatin regions via weak, intra- and intermolecular multivalent interactions (3, 5, 6, 9, 17, 75). While the molecular details underlying these interactions, whether occurring individually or with DNA, are being actively investigated, the influence of other inter-nucleosomal interactions on these phase-separated droplets remains largely unexplored. The ability of MBD proteins to undergo phase separation emerges as a compelling model for a general mechanism by which these (Figure 8) and potentially other chromatin-related proteins exert control over genome organization and transcriptional regulation (5, 19, 76, 77). To investigate this model, we focused on MBD2 and MBD3, two proteins significantly involved in modulating the transcriptional state of the genome. Our approach involves reconstituting a simplified microenvironment both *in vitro* and *in silico* to assess the LLPS of MBD2 and MBD3 and to identify the underlying molecular driving forces.

**Figure 8.**
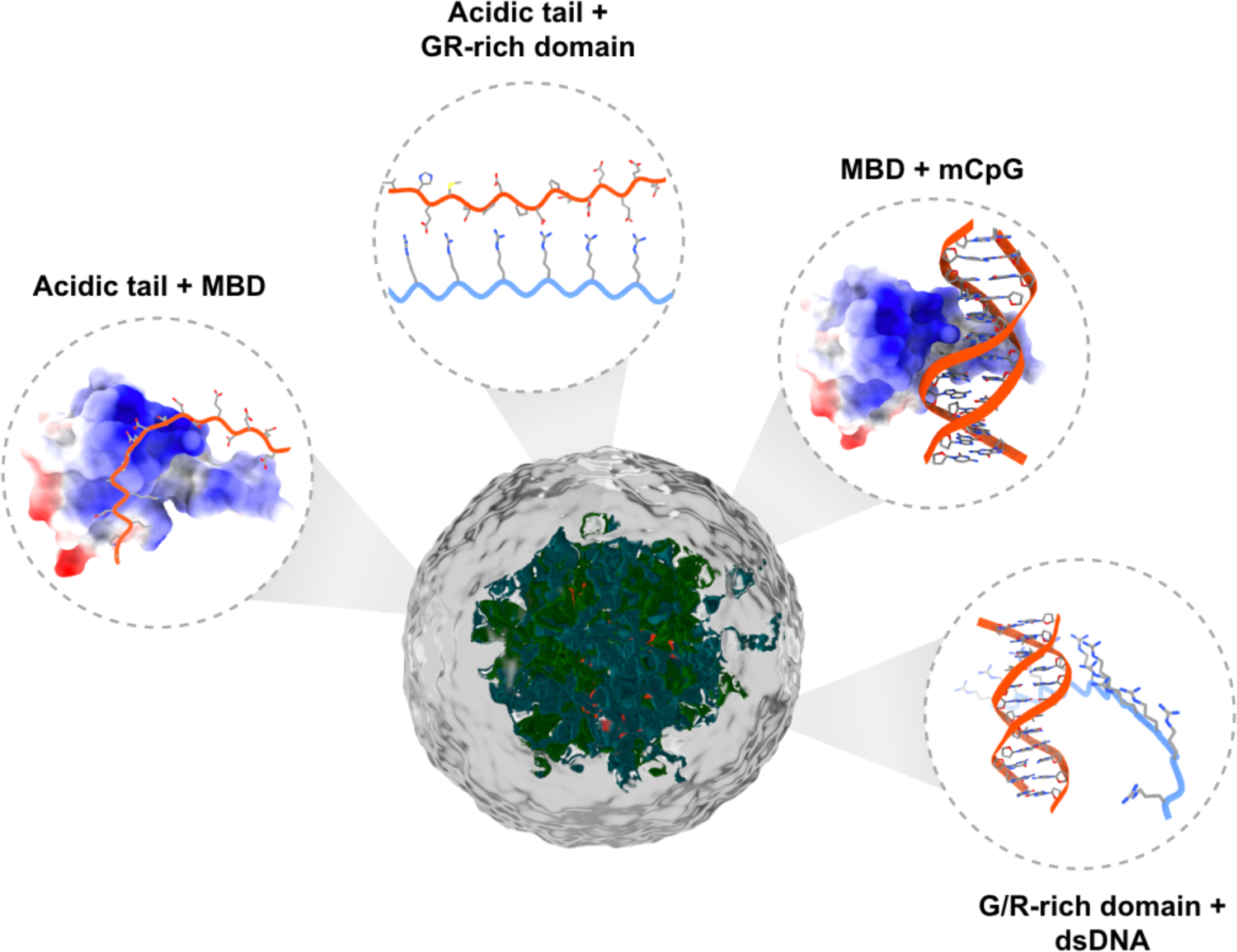
Model of the domain interactions governing MBD2/3 LLPS. The MBD is colored by the Coulombic surface (red = negative, white = neutral, and blue = positive).

Despite the inherent challenges of reconstituting large IDPs *in vitro*, we successfully established a robust expression and purification protocol for MBD2 and MBD3, enabling the production of significant quantities of both full-length and truncated proteins for characterization. Utilizing a combination of *in silico*, *in vitro*, and *in cellulo* approaches, we demonstrated the ability of MBD2 and MBD3 to undergo LLPS both individually and collectively, and in conjunction with (un)methylated DNA. Our findings demonstrate that the LLPS of these proteins is governed by a delicate balance of electrostatic and hydrophobic interactions, alongside a complex interplay of structural elements involving both protein-protein and protein-DNA interactions (Figure 8). Furthermore, MBD2 and MBD3 exhibit distinct phase behaviors due to their unique sequence properties, further modulated by temperature and protein concentration. These observations suggest the formation of large assemblies by MBD proteins, potentially serving as biological scaffolds for the recruitment of other factors. Notably, the addition of other components, such as the formation of complexes or the incorporation of methylated DNA, significantly enhances phase separation. By elucidating the interactions that drive the formation of MBD protein, we can gain valuable insights into the unique biochemical and cellular functions of heterochromatin in its condensed state. A crucial aspect of future research will involve deciphering how molecular activities are altered within condensates and understanding how these changes contribute to unique cellular functions in contrast to those observed in a dispersed state.

## Data Availability

Raw data and code are available upon reasonable request.

## Acknowledgments

We thank Dr. Michael Cosgrove for extending his expertise and providing critical feedback. We also thank Nathan McKean and Catherine Campbell for aiding with experimental work. We are grateful for the computational resources provided by Texas A&M High Performance Research Computing (HPRC).

