## Supplementary File for "Uncovering the molecular interactions underlying MBD2 and MBD3 phase separation"

### Supporting Information Figures

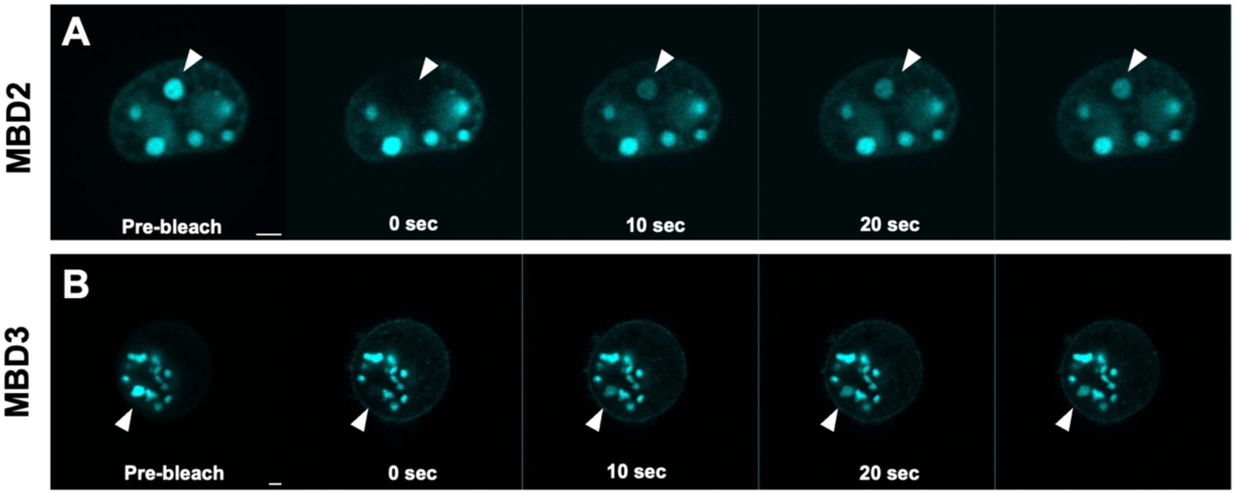


**Figure S1**. Pre- and post-fluorescence recovery after photobleaching (FRAP) images of live NIH-3T3 cells transfected with **A**. EGFP-tagged full-length MBD2 **B**. EGFP-tagged full-length MBD3. MBD2 and MBD3 droplet recovery is monitored from left to right as indicated by a white arrow within a 30-second time frame. Protein droplets were bleached using a 488 nm laser at 50% laser power. All cells are viewed at 100x using a Leica SP8 confocal microscope. The scale bar, represented by a white bar on the bottom right corner of the pre-bleach images, is 2 microns. Image montages were generated using FIJI/ImageJ.

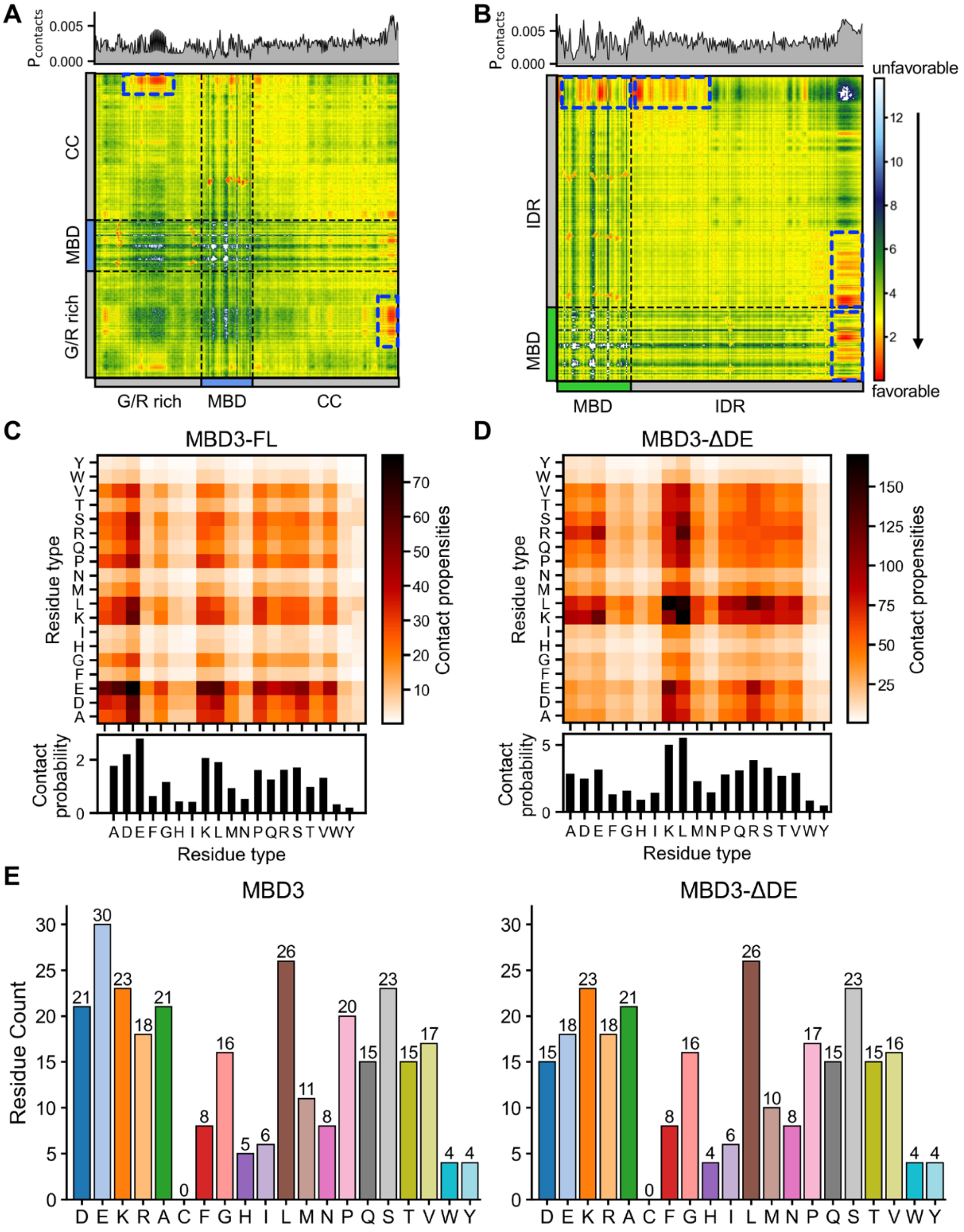


**Figure S2. A and B.** Intermolecular contacts of MBB2 and MBD3 proteins within the condensed phase. Preferential interactions are shown in red. The 1D contact map on the top is the average contact propensity per frame per residue. The CG coexistence simulations were conducted using the HPS-Urry model at 320K and 100 mM salt concentration. Blue boxes highlight dominant contact-prone regions. **C and D**. Intermolecular contact maps within the condensate between two residue types within the condensate of FL MBD3 and its ΔDE truncation, respectively. **E**. Bar charts of amino acid abundance in FL MBD3 and its ΔDE truncation.

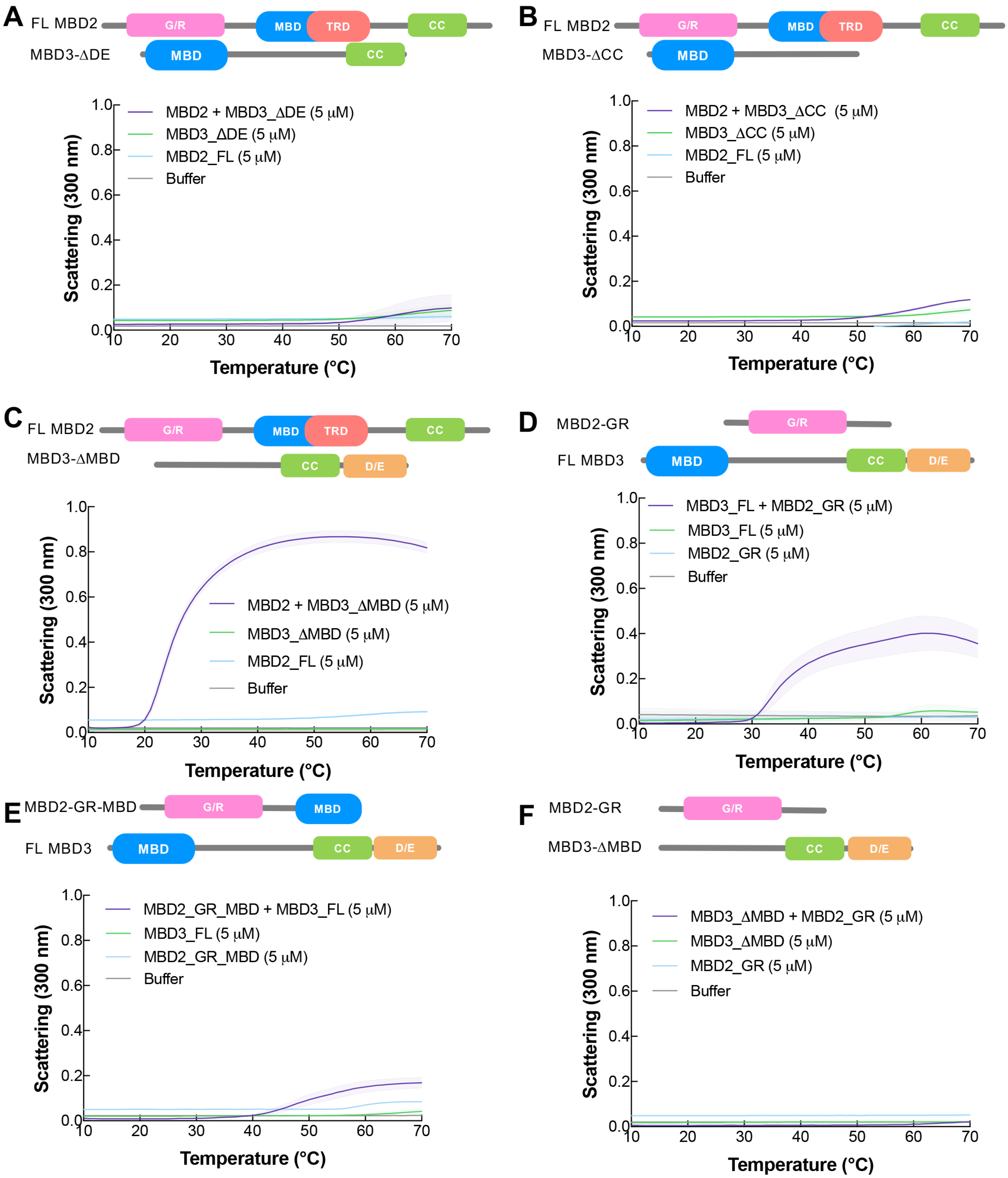


**Figure S3**. UV-Vis absorption spectra of **A**. FL MBD2 and ΔDE **B**. FL MBD2 and ΔCC **C**. FL MBD2 and ΔMBD **D**. FL MBD3 and GR **E**. FL MBD3 and GR-MBD **F**. GR and ΔMBD. The spectra show absorption as a function of the temperature of each protein individually (5 μM) and mixed together at an equimolar ratio (final concentration of 5 μM) in a phase separation buffer. The shading around each curve represents the standard deviation from the mean absorbance from technical replicates.

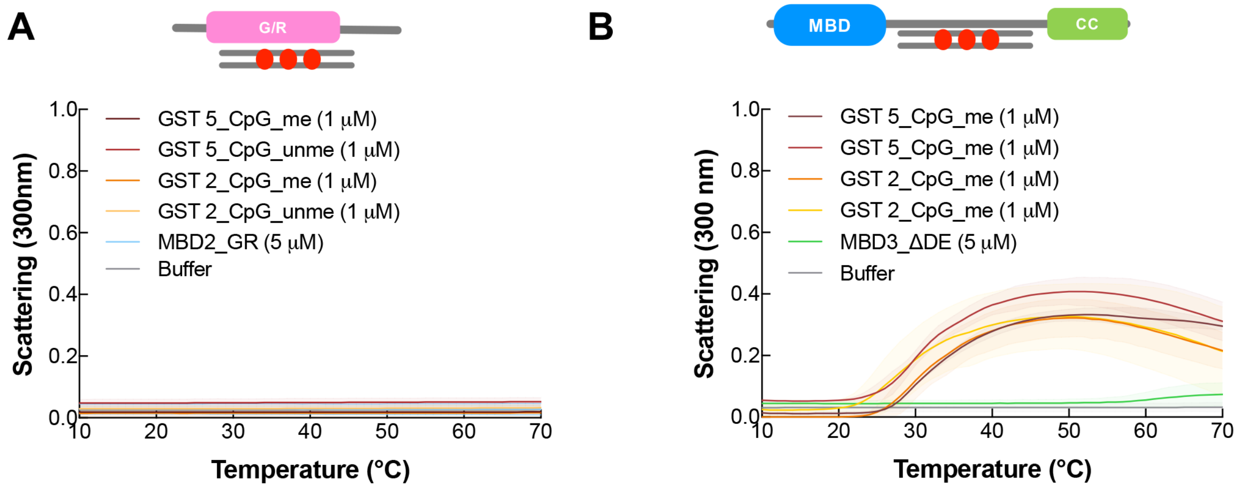


**Figure S4**. UV-Vis absorption spectra of **A**. GR and **B**. ΔDE individually and mixed with either unmethylated or methylated DNA that contains either 2 or 5 CpG sites in phase separation buffer as a function of temperature. Protein and DNA concentrations remained constant at 5 µM and 1 µM, respectively. The shading around each curve represents the standard deviation from the mean absorbance from technical replicates.

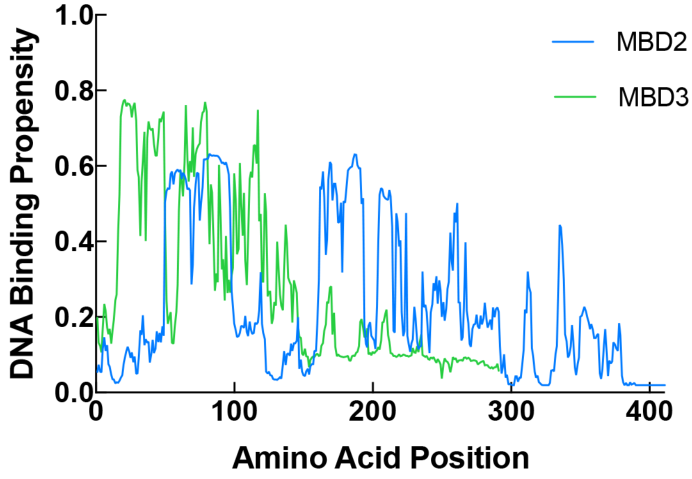


**Figure S5**. HybridDBRpred DNA binding propensity predictions for MBD2 and MBD3. A DNA binding propensity score above 0.28 indicates binding to DNA.
